# Morphology, geographic distribution, and host preferences are poor predictors of phylogenetic relatedness in the mistletoe genus *Viscum* L

**DOI:** 10.1101/433359

**Authors:** Karola Maul, Michael Krug, Daniel L. Nickrent, Kai F. Müller, Dietmar Quandt, Susann Wicke

## Abstract

Besides their alleged therapeutic effects, mistletoes of the genus *Viscum* L. (Viscaceae) are keystone species in many ecosystems across Europe, Africa, Asia and Australia because of their complex faunal interactions. We here reconstructed the evolutionary history of *Viscum* based on plastid and nuclear DNA sequence data. We obtained a highly resolved phylogenetic tree with ten well-supported clades, which we used to understand the spatio-temporal evolution of these aerial parasites and evaluate the contribution of reproductive switches and shifts in host ranges to their distribution and diversification. The genus *Viscum* originated in the early Eocene in Africa and appeared to have diversified mainly through geographic isolation, in several cases apparently coinciding with shifts in host preferences. During its evolution, switches in the reproductive mode from ancestral dioecy to monoecy imply an important role in the long-distance dispersal of the parasites from Africa to continental Asia and Australia. We also observed multiple cases of photosynthetic surface reduction (evolution of scale leaves) within the genus, probably indicative of increasing specialization associated with the parasitic lifestyle. Even compared with other parasitic angiosperms, where more host generalists than specialists exist, *Viscum* species are characterized by extraordinarily broad host ranges. Specialization on only a few hosts from a single family or order occurs rarely and is restricted mostly to very recently evolved lineages. The latter mostly derive from or are closely related to generalist parasites, implying that niche shifting to a new host represents an at least temporary evolutionary advantage in *Viscum*.

## 1. Introduction

Mistletoes are keystone resources in forests and woodlands because of their diverse interactions with the ecosystem’s fauna (Watson 2001). The mistletoe genus *Viscum* is well-known in Europe, Africa, and Asia since ancient times, where it served as fodder in Neolithic Europe (Heiss 2012). Especially *V. album*, the European or common mistletoe, is still valued as a medicinal plant for its alleged therapeutic activity in cancer and hypertension therapies (Deliorman et al. 2000; Kienle et al. 2009). Species such as *Viscum triflorum, V. tuberculatum*, and *V. album* represent food resources in different African and Asian countries (Bussmann 2006; Kunwar et al. 2005).

The genus *Viscum* (Viscaceae, Santalales) comprises from 70 (Wu et al. 2003) to 120 species (Nickrent 1997 onwards) distributed in the tropical and subtropical regions of Africa, Madagascar, Asia, and Australia, the temperate zones of Europe and Asia as well as the temperate southern Africa. One species, *V. album*, has been introduced and persists in the U.S. and Canada. All *Viscum* species are shrubby mistletoes, that is, obtaining water and nutrients via a multi-functional organ (haustorium) that penetrates the shoot of the host to connect with its vascular tissue. These parasites grow endophytically as cortical strands under the hosts’ bark (Kuijt 1969), unlike some species of Loranthaceae that form epicortical roots on the surface of the host branch. The genus *Viscum* is characterized by small, unisexual, insect and wind-pollinated flowers (Hatton 1965; Kay 1986) (Figure 1). The axillary or terminal inflorescences consist of petiolate or sessile cymes. Both dioecy, where individuals have either male or female flowers, and monoecy, where one individual carries flowers of both sexes, occur within the genus. Most African and Madagascan species are dioecious, whereas many Asian and all Australasian species exhibit monoecy (Barlow 1983a,b). The fruits are white, yellow, orange, or red typically one-seeded berries that are spread by birds. *Viscum* species retain the ability to photosynthesize to a greater or lesser extent. The leaves are either thick and leathery or scale leaves, often reduced to relic leaves of diminutive size. Furthermore, the genus includes an endoparasitic species, *V. minimum*, a South African endemic, that only emerges from the hosts’ tissue for reproduction.

**Figure 1.**
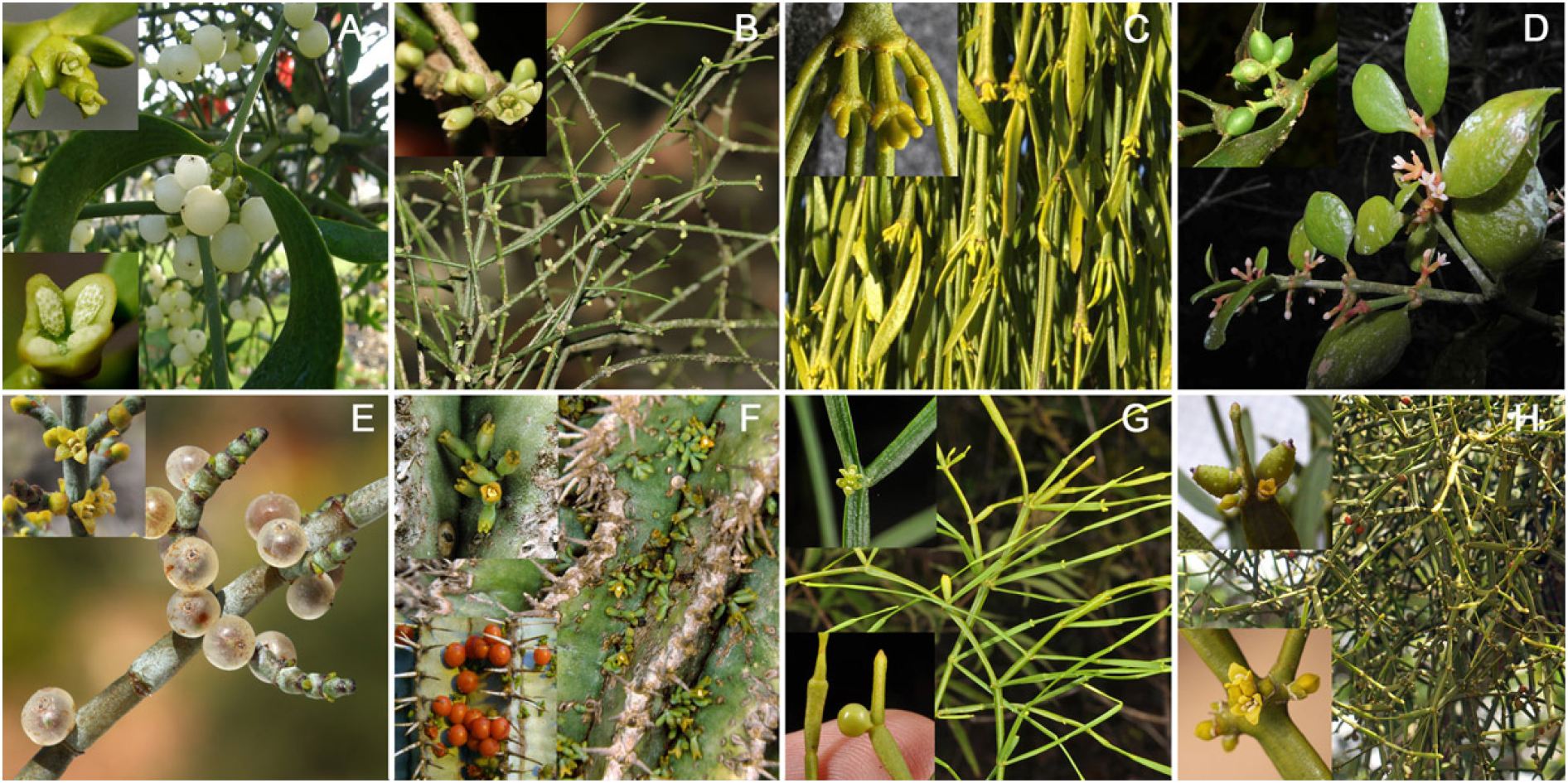
Growth forms of *Viscum*. **A.** *Viscum album* (clade B), a dioecious species from Europe. Plant in fruit. Insets: top – female flowers, bottom – male flower. Photo credits: Karola Maul, Gerhard Glatzel. **B**. *Viscum trachycarpum* (clade C), a dioecious, leafless species from Madagascar. Inset: male flowers. Photo credits: Peter Phillipson. **C**. Pendant shoots with young inflorescences of *Viscum whitei* (clade D), a monoecious species from Australia. Inset: closer view of inflorescences. Photo credits: Roger Fryer and Jill Newland. **D**. Flowering female plant of *Viscum cuneifolium* (clade E), a dioecious species from Madagascar. Inset: young fruits clustered in bibracteal cup. Photo credits: Peter Phillipson, Christopher Davidson (Flora of the World). **E**. Female plant with mature fruits of *Viscum capense* (clade F), a dioecious species from South Africa. Inset: male flowers. Photo credits: Marinda Koekemoer. **F**. Inflorescences of the monoecious, South African *Viscum minimum* (clade G) arising from the stem of a *Euphorbia* host plant. Insets: top – close-up view of male and female flowers, bottom – mature fruits. Photo credits: Karola Maul and Daniel Nickrent. **G**. *Viscum articulatum* (clade I), a monoecious, leafless species from Asia and Australia (here Philippines). Habit of mistletoe showing flattened branches. Insets: top – male flowers, bottom – fruit. Photo credits: Pieter Pelser. **H**. Branches bearing mature fruits of *Viscum combreticola* (clade J), a dioecious, leafless species from tropical east Africa. Insets: top – female flowers and developing fruits that are tuberculate when young, bottom – male flowers. Photo credits: Karola Maul, Bart Wursten. Clade names B—J (in parentheses) refer to our circumscriptions from phylogenetic inferences presented herein.

The patterns of host specificity within the genus differ widely. For example, *Viscum album* is known to parasitize more than 400 host species (Barney et al. 1998), whereas *Viscum minimum* grows only on two closely related species of *Euphorbia*. Besides this, several species such as *V. fischeri* occasionally parasitize other mistletoes (e.g. *Phragmanthera* or *Tapinanthus*), whereas some like *V. loranthicola* are obligate epiparasites on various species of Loranthaceae (Polhill and Wiens, 1998).

European mistletoes are well-known to biologists and the public, but the evolutionary history of *Viscum* is still elusive. Several morphological classification systems have been proposed for *Viscum* based on reproductive mode (monoecy or dioecy), the presence or absence of leaves, leaf shape, stem, and fruits, but also structures of the reproductive organs and inflorescences (summarized in Table 1). Although all African *Viscum* species have been monographed (Polhill and Wiens 1998) and their morphology analyzed cladistically (Kirkup et al. 2000), these infraspecific relationships remain to be tested with independent data. To date, the only molecular phylogenetic analysis of the genus was based on the nuclear large-subunit ribsosomal DNA (Mathiasen et al. 2008). Although this study sampled only 12 *Viscum* species and the phylogram lacked resolution, it did provide preliminary evidence of geographic rather than morphological clades. Expanding this preliminary molecular phylogenetic study to include more species and more genetic markers thus holds the potential to reconstruct the evolutionary history of *Viscum*.

**Table 1.** Summary of major classification systems for *Viscum*.

| Author(s) <sup>#</sup> | Basis for classification system | Number of species included (geographical distribution) |
| --- | --- | --- |
| Korthals (1839) | Presence vs. absence of leaves, monoecy vs. dioecy | 8 (Java, Sumatra, Borneo) |
| van Tieghem (1896) | Modification of Korthals' system, prioritizing the composition and position of inflorescences | 27 (Madagascar, continental Africa, Asia, Europe) |
| Engler and Krause (1935) | Inflorescence characters: position, number of flowers; phyllotaxy | 42 (continental Africa, Madagascar, Europe, Asia) |
| Danser (1941) | Structure of inflorescences | 24 (India, SE-Asia, Malaysia, Australia) |
| Balle (1960) | Structure and composition of inflorescences | 99 (Madagascar, continental Africa, Europe, Asia, Oceania) |
| Polhill and Wiens (1998) | Macroscopic morphological characteristics of the whole plants | 45 (continental Africa) |

<sup>#</sup> Several other regional treatments exist (e.g., Rao 1957, Sanjai and Balakrishnan 2006, or Barlow 1997), which we here omitted for simplicity.

Here, we use 33 sequences of the plastid marker *rbcL*, *37 of trnL-F*, and 19 of matK as well as 110 sequences of the nuclear ribosomal internal transcribed spacers 1 and 2 (ITS-1, –2) plus 21 sequences of the 18S ribosomal RNA gene from species and subspecies of *Viscum* and several outgroup taxa of Santalales to elucidate the diversification pattern of these important keystone plants. Our final phylogenetic tree of 110 taxa was subjected to extensive phylogenetic analyses employing maximum likelihood and Bayesian inference. We focused specifically on analyzing whether cladogenesis correlates primarily with morphological differentiation, geographic range distribution, or host preference. To this end, we estimated the age of the genus and the infrageneric divergence times using a relaxed molecular clock. These data provided a solid basis to reconstruct the geographical origin of *Viscum* and trace its subsequent distributional history across four continents. We also reconstructed by maximum likelihood the evolution of morphological characters such as foliage type and reproductive mode, considering phylogenetic uncertainty. We mapped and reconstructed the host preferences and the evolution of host ranges to test if diversification in *Viscum* includes significant host range shifts. This study will aid in understanding diversification of parasitic plants that are important components of flora-faunal interactions in many ecosystems.

## 2. Materials and Methods

### 2.1 Taxon Sampling

The classification of Santalales families used here follows Nickrent et al. (2010). We used 220 sequences of 5 nuclear and plastid markers from 59 *Viscum* taxa (incl. 2 subspecies) and, in total, 73 outgroup taxa from Viscaceae and three other Santalales families for the final computation of a 110-taxa phylogeny. Our data sets included sequences generated in this study (from one plant individual each), as well as additional data from Genbank. Supplemental Tables S1 and S2 detail the origin and voucher information for all taxa included here. Despite the low availability of useful *Viscum* specimens in herbaria, we successfully sampled the genus representatively in terms of geographical distribution, and morphological trait variability (see 2.3 and 2.4 below).

### 2.2 Experimental procedures

We sequenced the nuclear ribosomal ITS region (internal transcribed spacers 1, 2 and 5.8S rDNA) for 59 *Viscum* taxa and several outgroups within and outside Viscaceae. In addition, we sequenced the plastid loci *rbcL* and *trnL-F* for nine outgroup and *Viscum* species. DNA was isolated either from herbarium specimens or from fresh or silica-dried tissues using the NucleoSpin Plant II Kit (Macherey-Nagel) according to the manufacturer’s protocol. PCR amplification was performed in a 25 μl reaction mix containing 1 μl DNA template (10–30 ng/μl), 2 mM of MgCl_2_, 0.2 mM of each dNTP, 0.8 μM of each primer, 1 U GoTaq Flexi DNA polymerase (Promega) and 1× GoTaq Flexi Buffer. The PCR program consisted of 5 min at 94°C, 35 cycles of each 1 min at 94°C, 1 min at 48°C, and 0.45 min at 68 °C, plus a final extension step of 10 min at 72 °C.

In most cases the amplification of the ITS region was successful with the ITS4 and ITS5 primers (White et al. 1990). We used the following newly designed, *Viscum*-specific primers: SEQITS2_VISCUM (5’-AACGACTCTCGRCAATGG-3’) with ITS4 and SEQITS1_VISCUM (5’-TTGCGTTCAAAAACTCAATGA-3’) with ITS5 to amplify the region in two overlapping halves when only low-quality or fragmented template DNA was available. We amplified *rbcL* and *trnL-F* using the rbcL-1F primer (Olmstead et al. 1992) in combination with *rbcL*-1368R (Fritsch et al. 2001), and the *trnL-F* universal primers (Noben et al., 2017), respectively. DNA sequencing was carried out by GATC Biotech (Germany) or Macrogen Inc. (Netherlands). We complemented our dataset with several taxa deposited in Genbank (Table S1) to construct two final data sets: data set 1 consisted of the nuclear 18S rRNA gene concatenated with the plastid markers *matK, trnL-F*, and *rbcL* (Supplemental Table S1); data set 2 consisted only of the entire ITS region (Supplemental Table S2).

### 2.3 Phylogenetic analyses

Sequence data editing and manual alignment were performed using PhyDE v0.9971 (http://www.phyde.de). Because of high variability in the nuclear ribosomal internal tran-scribed spacers across genera and families (51.6% pairwise identity over all taxa; pairwise identity within *Viscum*: 72.4%), we applied a two-step approach, which consisted of the reconstruction of a backbone guide tree to aid the resolution of deep nodes in the phylogenetic tree based on a multi-marker data set with the more slowly evolving nuclear 18S rRNA gene and the plastid markers (data set 1) and a subsequent inference of lower-level relationships on a condensed but near-complete matrix of the faster evolving ITS region (data set 2) using constraints resulting from step 1. Rather slowly evolving markers have a higher chance to resolve deep nodes more accurately and suffer less from homoplasy, amongst others (e.g., Wicke and Schneeweiss 2015).

We aligned the data set 1 manually and excluded mutational hotspots in the *trnL-F* spacer where homology assessment was ambiguous (all alignments available from *Mendeley Data*). The backbone guide tree was computed using Bayesian inference (see below), for which we added information from simple gap coding (hereafter: SIC) (Simmons and Ochoterena 2000), obtained with *SeqState* 1.4 (Müller 2005), thereby appending another 266 characters to our data matrix.

Additionally, we generated ITS region alignments for the following subgroups within which the variability of this nuclear marker region still allowed confident homology assessment: 1) *Arceuthobium*, 2) *Korthalsella* and *Phoradendron/Dendrophthora*, 3) Loranthaceae, 4) Santalaceae incl. Amphorogynaceae, 5) *Notothixos*, and 6) *Viscum*, with the *Phoradendron/Dendrophthora* clade as outgroup. These data subsets were aligned manually (*Viscum*) or using *PRANK* v1.3 (Löytynoja and Goldman 2005; default settings) to generate group-wise sub alignments, to be used as anchors. Phylogenetic relationships within these specific data sets were computed by Bayesian inference, and the reconstruction of a full phylogeny was then constrained with the backbone tree through genera- or cladewise addition. To obtain a complete ITS region alignment of 110 taxa, some of which having served already as sub-alignment anchors, the resulting tree (Supporting Figure S1) was used as a guide tree for *PRANK* under default settings (data set 2; Supporting Table S2).

The final ITS region alignment was subsequently analyzed with and without simple indel coding using Bayesian inference (BI) and maximum likelihood (ML), respectively. Bayesian analyses were conducted with *MrBayes* v3.2.5, x64 MPI version (Huelsenbeck and Ronquist 2001) under the GTR+Γ model. We analyzed six runs with four parallel chains, each with one million generations for the family-specific data sets and ten million generations for the combined analysis, allowing for a burn-in phase of the first 25% of all iterations. ML inferences were computed with RAxML v8.1.2 under the GTR+Γ model and 10,000 bootstrap replicates (Stamatakis 2014). To evaluate the robustness of our data set 2 alignment and our approach in general, we also employed *Guidance II* (Sela et al. 2015) in combination with *mafft* (Katoh et al. 2002) but in the absence of a guide tree and under three different stringency settings, leading to the elimination of differing proportions of uncertain or homoplasious sites (optimized alignment variants available from *Mendeley Data*). The resulting three data sets were used to re-compute phylogenetic trees with RAxML and MrBayes as described above.

Results and trees were visualized in *Treegraph* 2.4 beta (Stöver and Müller 2010), or with the R statistical computing framework in combination with the *R* packages *ape* (Paradis et al. 2004) and *phytools* (Revell 2012).

### 2.4 Molecular clock dating and estimation of the distribution range evolution

We performed a molecular dating analysis using *PhyloBayes* v3.3 (Lartillot et al. 2009) with the CAT Dirichlet process mixture for among-site substitution heterogeneities (Lartillot 2004) and CIR (Lepage et al. 2007) as clock relaxation process in addition to a log-normal autocorrelated relaxed clock (LN) analysis. Using our combined data set 2 and the results from the Bayesian and ML inferences with and without SIC, we applied a primary calibration to constrain our analyses with fossil data with minimal ages according to the records’ upper bounds as follows: Loranthaceae – 51 million years (mya; Macphail et al. 2012), Santalaceae – 65 mya (Darrah 1939; Christopher 1979), *Arceuthobium* – 52 mya (Krutzsch 1962), and the most recent common ancestor of *Viscum* to 28 mya (Mai et al. 2001); no root age constraint was set. Per analysis, we ran two parallel chains from which we sampled every 5^th^ generation until 20,000 were collected. MCMC chain convergence was assessed via the discrepancy between the posterior averages obtained from independent runs and the effective size of several summary statistics. Because of the good convergence of both chains, we merged the chains and computed the consensus divergence age estimates per every input tree. Trees were visualized using the *ape* package in R (Paradis et al. 2004).

*LaGrange* v2 (Ree and Smith 2008) was run to reconstruct the evolution of geographic ranges in *Viscum* using the four phylogenetic trees (BI, BI-SIC, ML, ML-SIC) and the corresponding *PhyloBayes* inferred root ages (BI: 142.02 mya (CIR), 120.258 mya (LN), BI-SIC: 144.182 mya (CIR), 122.172 mya (LN), ML: 140.969 mya (CIR), 121.558 mya (LN), ML-SIC: 144.195 mya (CIR), 118.287 mya (LN)). For the analysis, we classified seven relevant regions: Europe, Africa north of the Sahara, Sub-Saharan Africa, Madagascar and Comores, continental Asia (north of the Wallace line), Australasia (south of the Wallace line), and the Americas. Distribution ranges of the extant species were coded as a binary text file indicating the presence or absence in these regions (Table S3). We input the species range data matrix, and the phylogenetic tree into the *LaGrange* configurator adding the root node ages from our molecular dating analysis. We excluded the following direct transition combinations from the adjacency matrix: Madagascar and Americas, Europe and Madagascar, and Europe and Australasia, respectively, because we considered such transitions as highly unlikely. All rate parameters were estimated with dispersal constraints consistently set to 1.0, and we calculated with only one-time period from 0 to the respective root age. We visualized the results with *Treegraph* 2.4 beta (Stöver and Müller 2010).

### 2.5 Ancestral state estimation

Information on the morphological characters regarding foliage was coded with three states as 0 = leafy, 1 = scale leaves, and 2 = leaves and scale leaves; the plant reproductive mode was coded as 0 = monoecious, 1 = dioecious, and 2 = bisexual flowers (Table S4; available from *Mendeley Data*). Ancestral state reconstruction was conducted for both features separately with *BayesTraits* v2.0 under the MultiState option with 1000 maximum likelihood attempts (Pagel et al. 2004). To test all possible topologies of the ingroup, we performed this analysis based on the BI trees with and without SIC, and the ML tree with SIC, respectively (Supporting Figure S2). We did not consider the ML tree without SIC because the topology was identical to the BI tree without SIC.

Host range distribution within the genus *Viscum* was primarily assessed on the basis of an extensive literature research (Table S5; also available from *Mendeley Data*). To evaluate whether a broad host range is a derived character, we traced the evolution of host ranges by inferring the ancestral number of potential host species or host genera across the consensus topology via maximum likelihood ancestral state reconstruction, implemented in the *R* packages *ape* (Paradis et al. 2004) and *phytools* (Revell 2012).

## 3. Results

### 3.1 Phylogenetic relationships

Maximum likelihood and Bayesian inference based on the concatenated nuclear and plastid gene dataset provided a first comprehensive and statistically well-supported phylogenetic tree of the mistletoe genus *Viscum* (Fig. 2 and Supporting Figures S1, S2, S3; original tree files in *newick* format available from *Mendeley Data*). Our analyses resolve *Notothixos* as sister to all other Viscaceae and Phoradendreae as sister to *Viscum*. Within *Viscum*, we can define ten well-supported clades: Clade A consists of three species from eastern Africa, whereas species of clade B occur in sub-Saharan Africa (*V. congolense*), Northern Africa, Southern Europe, and the Near East (*V. cruciatum*), temperate continental Asia (*V. nudum and V. coloratum*), and *Viscum album*, which is widespread from England to Japan. Madagascan species cluster in both clade C, which exclusively comprises Madagascan species, and clade E, that also contains one species occurring also in continental Africa (*V. decurrens*) and one species known from continental Africa and the Comoros (*V. triflorum*). Early diverging from clade E, although with low support, is clade D consisting of endemic Australian species (*V. whitei, V. bancroftii*). Clades F, G, and H contain species with rather small geographic ranges in Southern Africa, except for *V. tuberculatum*, which expands into eastern Africa. While clade I contains species mostly occurring in both continental Asia and Australasia, Clade J consists of species from sub-Saharan Africa.

**Figure 2.**
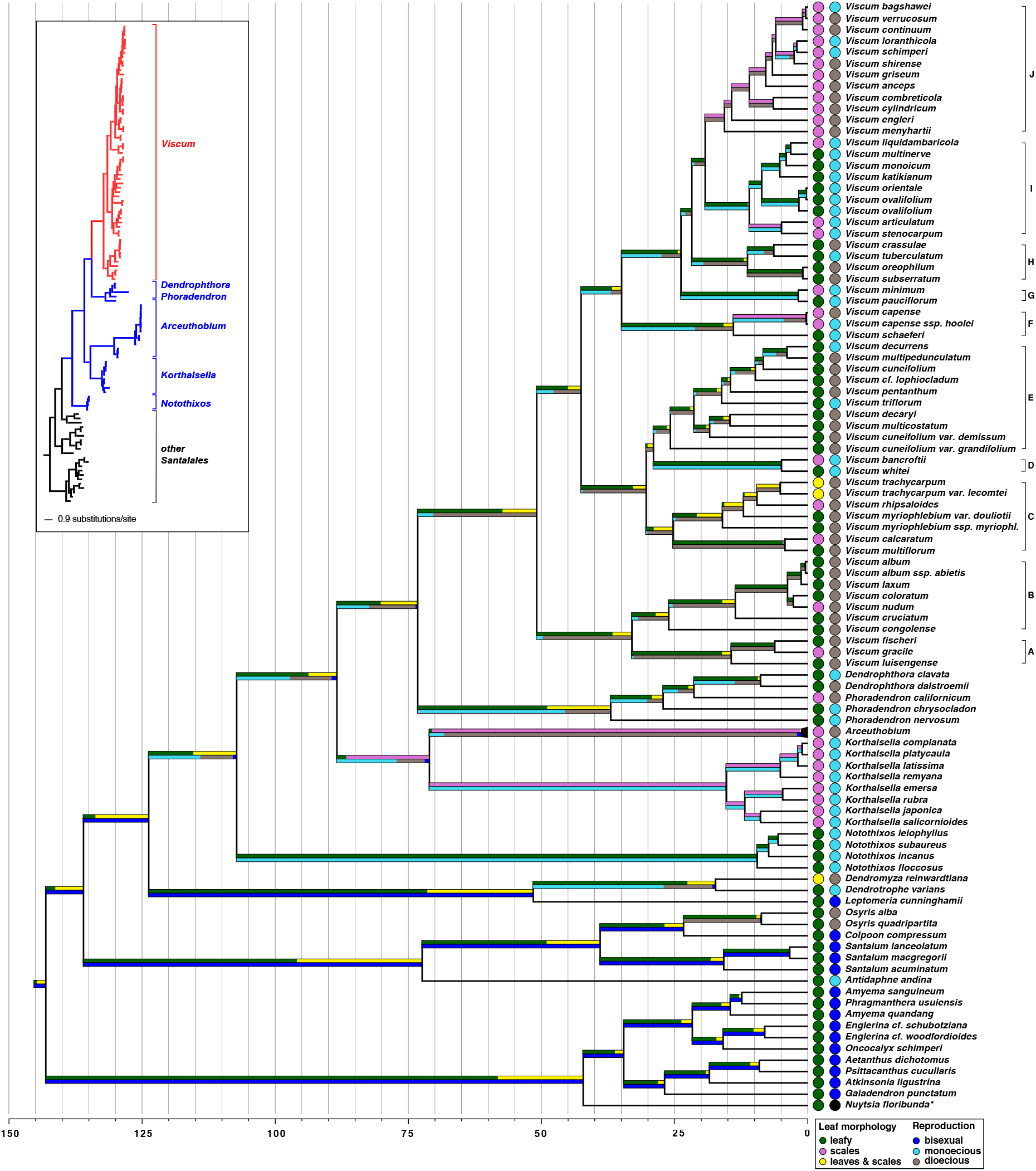
Evolution of leaves and reproductive systems in *Viscum*. The ancestral states of foliage and reproductive systems are illustrated by colored stacked bars below or above branches of the BI-SIC tree, obtained from the analysis of the ITS region alignment (data set 2), respectively. The lengths of the individual stacks represent, proportionally, the probability of the ancestor adopting a certain state (color-coded as detailed in the bottom right corner). The lengths of the stacks (corresponding to 100 % cumulated probability per branch) are scaled according to evolutionary time in millions of years. Colored circles at the tips of the tree show the extant foliage or reproductive morphology per species, which was used for ancestral state estimation. The inset depicts the simplified phylogram of the same MB-SIC tree, which is provided in full in Supporting Figure S2-A. *Note that the category „polygamous“(more specifically, polygamonoecious) was not included here, which is the actual reproductive condition in *Nuytsia*.

Our analysis showed that some taxa are not monophyletic: *V. myriophlebium* subsp*. myriophlebium*/*V. myriophlebium var. douliotii, V. cuneifolium*/*V. cuneifolium var. grandifolium*/*V. cuneifolium var. demissum, Viscum ovalifolium*. The ITS sequences of both specimens of *V. ovalifolium* and *V. orientale* are like one another, forming a clearly definable group.

Minor topological differences between our various phylogenetic reconstruction methods (BI with and without SIC, and ML with and without SIC, respectively) only occur within two outgroup clades (Loranthaceae and *Notothixos*), and on three positions inside *Viscum*. Both trees obtained with SIC (BI and ML) are congruent within *Viscum*, except for the topology of clade I: In the ML-SIC tree, the *V. ovalifolium/V. orientale* subclade is sister to the remaining clade, whereas the *V. arcticulatum/V. stenocarpum* subclade is sister to the remaining clade I in all other trees. No incongruency is seen between inferences without SIC (Fig. S3). Within *Viscum* they differ slightly from the SIC trees in the position of *V. menyhartii* in clade J and in the position of *V. minimum* and *V. pauciflorum*, which is sister to clade H with low support in the trees without SIC but forms a well-supported discrete clade being sister to clades H-J in the SIC analyses (Fig. S3). Except for the branch leading to extant *Arceuthobium* species, phylograms (Supporting Figures S2 and S5) from any of the conducted analyses show no conspicuously long branches within or between clades that could hint to a long-branch attraction phenomenon or elevated nucleotide substitution rates. Also, optimization of the alignment of data set 2 has no influence on the recovered topology (Supporting Fig. S4; results also available in *newick* format from *Mendeley Data*), only affecting node support values. These results suggest that the primary nucleotide data already exhibits a rather powerful phylogenetic signal for a robust phylogenetic inference even without SIC.

### 3.2 Molecular dating and ancestral range distribution

Our *PhyloBayes*-based divergence time estimations indicate that Viscaceae diverged in the late Lower Cretaceous, with an origin between 124.72 ± 31.55 mya (CIR) and 102.15 ± 33.67 mya (LN) for the stem group and 108.06 ± 27.35 mya (CIR) and 86.01 ± 27.94 mya (LN) for the crown group (Fig. 2 and Table S5). The root node age estimates based on our BI-SIC tree vary between 144.18 ±36.96 mya (CIR model) and 122.17 ± 41.86 mya (LN model. The reconstruction of the ancestral distribution suggests an Australasian origin of Viscaceae (more than two log-likelihood units in all analyses) (Fig. 3; Table S6). The genus *Viscum* separated from Phoradendreae 73.78 ± 18.99 (CIR), or 61.76 ± 19.62 (LN) mya. The crown group of *Viscum* evolved in the early Eocene (CIR: 51.31 ± 13.49 mya, LN: 45.61 ± 14.48 mya), when the ancestor of clade A/B diverged from that of the remaining *Viscum*.

**Figure 3.**
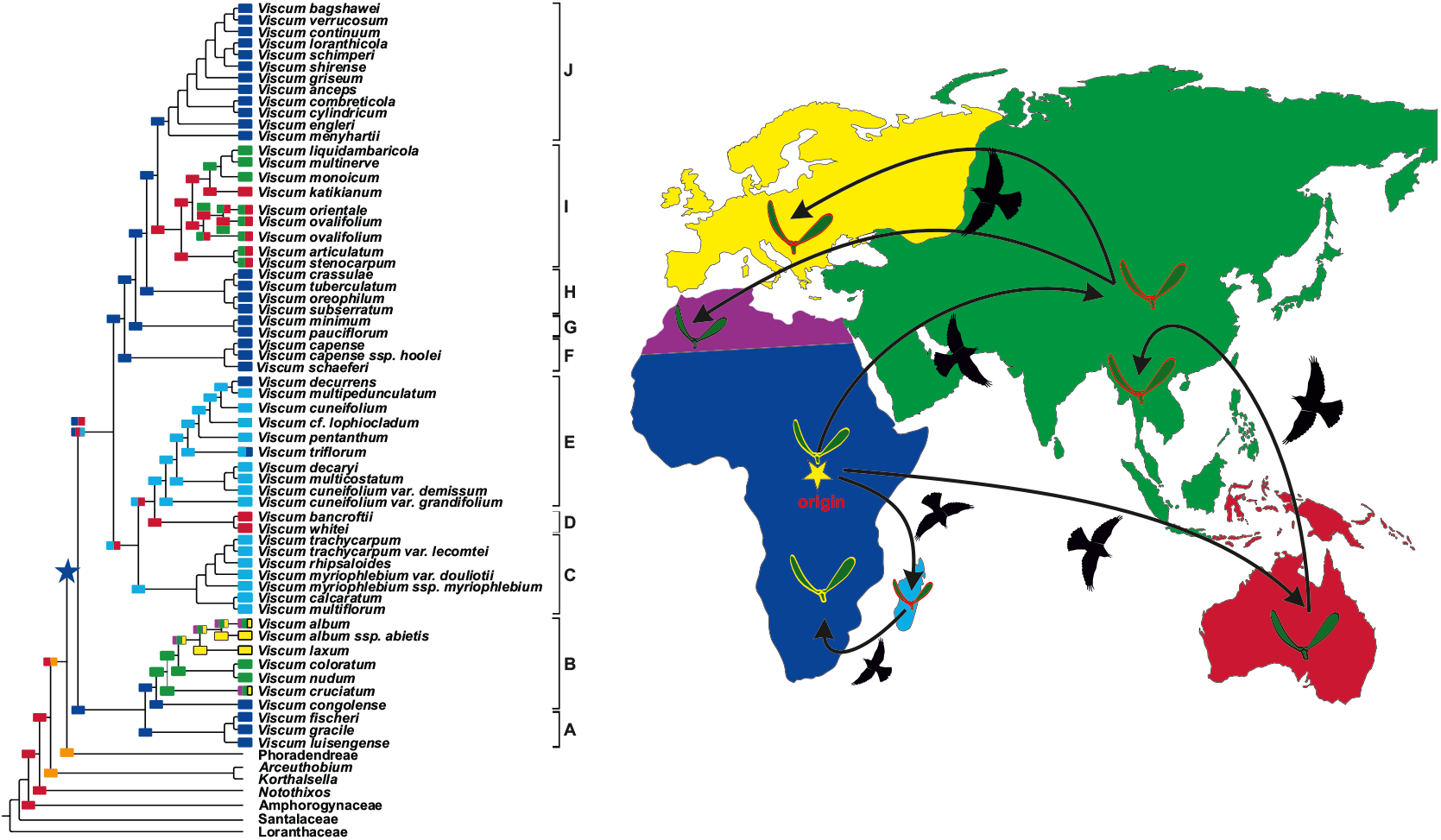
Biogeographical history of *Viscum*. The most probable ancestral geographic ranges as well as the extant ranges are illustrated by colored squares at all nodes on the dated phylogenetic BI-SIC tree of *Viscum* and outgroups, based on data set 2 (ITS region). Two different squares on top of each other indicate an ambiguity in the reconstructed geographic ranges, where two or more ancestral areas were equally likely (see Table S6 for details). Distributions exceeding one geographic region are shown by multi-color squares. The colonization history of Australia, continental Asia and Europe from Africa is graphically summarized. An arrow indicates the independent colonization event, and the star suggests the likely origin of *Viscum* in Africa.

The Lagrange results do not clarify the geographical origin of *Viscum*, although Africa as the ancestral area of the *Viscum* stem group has slightly better support than alternatives. The African range is retained throughout the whole backbone of *Viscum*. Continental Asia was most likely colonized 29.72 ± 10.19 (LN), or 26.38 ± 7.85 (CIR) mya (clade B). The Madagascan clades C and E, together with the Australasian clade D separated from the ancestor of remaining *Viscum* taxa 42.86 ± 11.24 (CIR), or 40.9 ± 12.95 (LN) mya. Given the tree topology is correct, Africa seems to have been colonized twice from Madagascar (CIR: 16.36 ± 4.82, LN: 16.6 ± 5.65 mya; CIR: 4.06 ± 1.55, LN: 3.27 ± 1.43 my). *Viscum* recolonized Australasia most probably 24.27±7.78 (LN), 19.47± 5.45(CIR) mya from Africa at the base of clade I (*V. articulatum*). Slightly higher support values suggest that *Viscum* spread to continental Asia three times independently from Australasia within Clade I (LN: 9.76 ± 3.7, CIR: 5.36 ± 1.88 mya; LN: 7.04 ± 2.89, CIR: 5.11 ± 1.99 mya; LN: 3.35 ± 1.84, CIR: 1.76 ± 0.97 mya; results from alternative tree topologies: Supporting Table S6).

### 3.3 Ancestral state estimation

The reconstruction of reproductive modes suggests that dioecy likely was the ancestral sexual condition in *Viscum* (posterior probability [pp]: 0.81 for dioecy versus 0.19 for monoecy) (Fig. 2; Supporting Fig. S6 and Table S7 for alternative tree topologies). Clades A, B, and C contain only dioecious species. Within the remaining clades reproductive mode varies. Monoecy evolved at least eight times independently, being replaced again by dioecy in five cases. Our analysis suggests that the most-recent common ancestors of the (Austral) Asian clades D and I likely both were monoecious (pp: 1.00 (D), 1.00 (I); Fig. 2; Fig. S6 and Table S7).

Our ancestral state reconstruction of foliage evolution suggests that the *Viscum* ancestor was leafy (pp: 0.72). The reduction of leaves occurred independently ten times during the evolution of *Viscum*, nine times from an ancestrally leafy condition and once from an ancestor with both leaves and scale leaves (Fig. 2). The extreme reduction to minute scale leaves evolved rather late during the diversification of *Viscum*, with one exception: within the African clade J, scale leaves are present in all species, thus it is likely that this feature was already present in the ancestor of this clade. Our analyses suggest that full foliage was maintained over ca. 19.61 mya (LN) or 25.71 mya (CIR), after the reconstructed origin of the genus (see below), and reject the possibility of a recurrence of the leafy habit in a clade that had previously already evolved scale leaves. Conversely, our reconstruction suggests that within clade C, leafiness has evolved again from the intermediate habit of both leaves and scales, once from the clade’s most recent common ancestor and a second time within the clade.

Visual inspection of host range distribution within the genus suggests that a shift in host specificity may have contributed to speciation and diversification in *Viscum* (Fig. 4). Based on an extensive literature search (summarized in Supporting Table S5), we observe that several sister taxa show distinct host specificities (e.g. *Viscum album* subsp. *album* and *V. album* subsp*. abies*; *V. pauciflorum* and *V. minimum*; the *V. schimperi*/*V. loranthicola*/*V. shirense* clade). Many species grow on core rosids. A clear ancestral order cannot be identified, partly because of the lack of host information for some taxa. The number of potential hosts is highest in *V. album* subsp*. album* and covers the broadest taxonomic diversity, ranging from basal angiosperms to lamiids and asterids (Fig. 4; Supporting Table S5). A few other Asian taxa such as *V. articulatum* are known to parasitize many different host plants, too, but the diversity of their hosts is narrower than that of *V. album* subsp*. album*.

**Figure 4.**
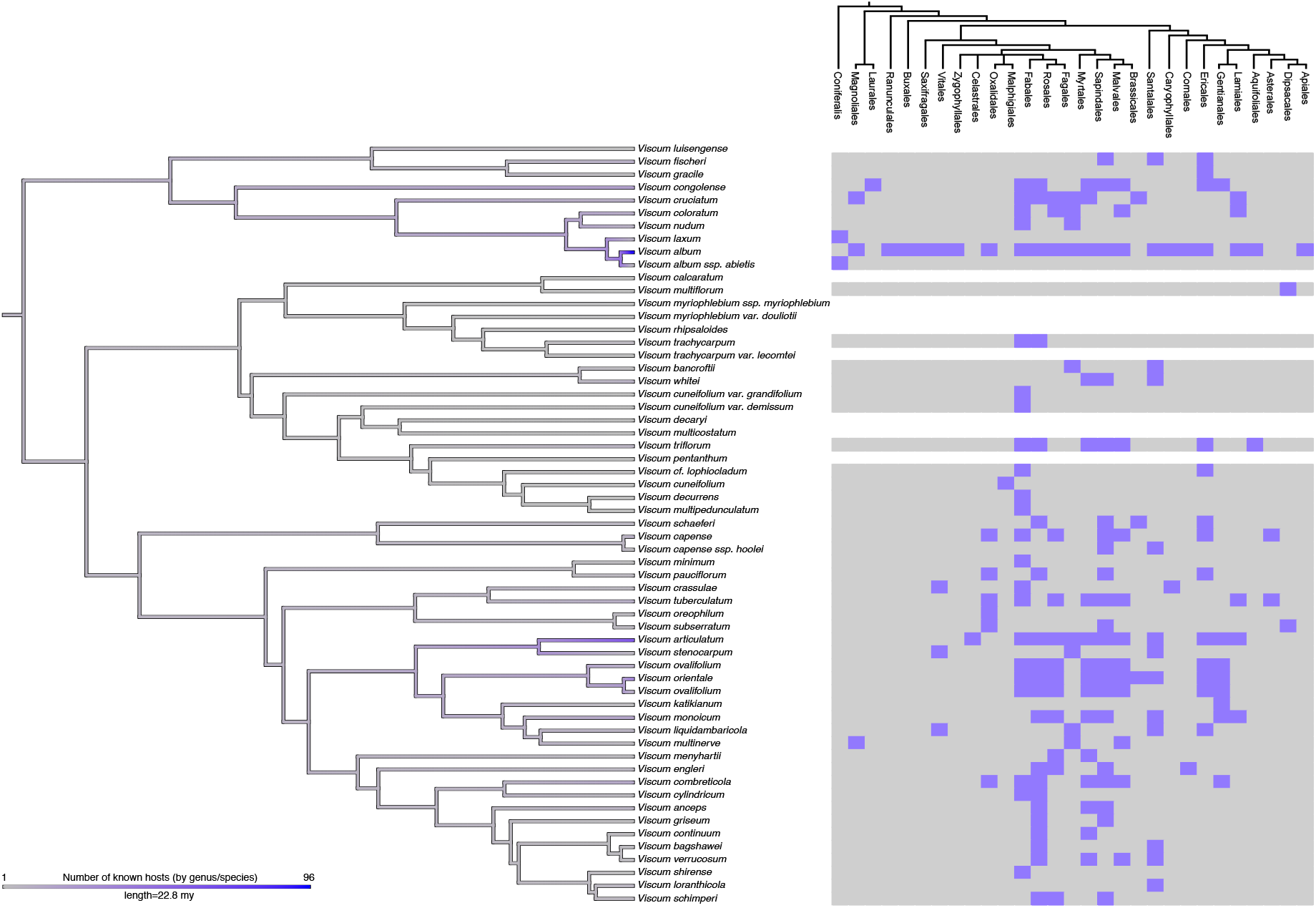
Host range distribution in *Viscum*. The number of ancestral hosts as reconstructed by maximum likelihood analysis of the number of described hosts (by family and genus, where available) is illustrated by a color gradient across the BI-SIC tree, obtained from data set 2. The number of plants accepted as hosts is shown with a grey (small) to blue (large) color gradient. The systematic position (by order) of the preferred hosts per species is indicated on the right-hand side, where the ability to parasitize species of a given order is shown in blue; gray – no hosts known from that order; white – no host information available.

## 4. Discussion

### 4.1 Diversification of *Viscum* is geography-driven – to some extent

Using nuclear and plastid markers, we here showed that cladogenesis within the mistletoe genus *Viscum* is more consistent with geographical ranges of the species they contain than with classifications based on morphology. Although some geographic patterning exists, thus corroborating to some degree an earlier hypothesis of a geography-driven diversification pattern (Mathiasen et al. 2008), the geographic distribution pattern of the extant *Viscum* species is complex and cannot alone be used to conclude interspecific relationships within the genus. The independent colonizations of Europe, continental Asia, as well as the recolonizations of the African continent and Australasia does not allow the discrimination of clades based solely on their distribution. We found that African *Viscum* species are polyphyletic (Figs. 2 and 3), present in six different clades. We also present evidence that the Australasian-continental Asian clade I most likely split from the African species possibly by stepping stone dispersal, although we cannot exclude that Australasia has been colonized via continental Asia (see below). In clade B, the species of temperate Asia appear to be more closely related to those occurring in temperate to subtropical regions of Europe, the Middle East, the Mediterranean and North-Saharan Africa. All other Australasian/Continental Asian species are phylogenetically distinct (clade I). For two species, where subspecific taxa were sampled (*V. cuneifolium* and *V. myriophlebium*), the accessions were not monophyletic at the species level. Future research using a greater sampling density with multiple accessions per species will be necessary to address the potential paraphyly of those taxa and to clarify the taxonomy within the genus *Viscum*. The estimation of species numbers in the genus has a broad range (see 1; Nickrent 1997, and onwards; Wu et al. 2003;), and our study thus covers between 50 % and 80 % of the recognized *Viscum* diversity. Although we consider our inference a valuable and rather robust first hypothesis of the evolutionary history of *Viscum*, analyses with a more exhaustive species representation might yield higher statistical support of the ancestral area and trait estimations, and could contribute towards the clarification of evolutionary processes of *Viscum*. Including multiple specimens identified as the same species should also be included in future work, especially to address the herein reported paraphyly of some taxa.

Besides our detailed analysis of the genus *Viscum*, the topology of the phylogenetic tree based on our concatenated four-marker “backbone” data set (Fig. S1) is congruent with a previously published phylogeny (Der and Nickrent 2008) regarding all deeper splits of the sandalwood order, Santalales. Our analyses of nuclear and plastid markers strongly support the monophyly of Viscaceae, in line with earlier studies (Soltis et al. 2000; Der and Nickrent 2008; Vidal-Russell and Nickrent 2008). Our multigene data set resolves *Notothixos* as sister to the remaining Viscaceae genera, and *Dendrophthora/Phoradendron* as sister to *Viscum*, which contradicts earlier data (Wiens and Barlow 1971; Der and Nickrent 2008; Mathiasen et al. 2008) and prompting for further studies, ideally involving phylogenomic approaches.

### 4.2 Eocene origin of *Viscum* results in the subsequent dispersal of a widely distributed ancestor

Our analyses based on Bayesian relaxed clock approaches date the crown group estimates of Viscaceae to 108.06–84.47 mya (Fig. 2; Supporting Fig. S6 and Table S7) and the reconstruction of the ancestral geographic range indicates an Australasian distribution. A Gondwanan origin of the family agrees with Barlow (1990), rejecting the hypothesized Laurasian origin suggested by Wiens and Barlow (1971). After the split of *Notothixos*, the ancestor of the remaining genera dispersed to the Americas during the Upper Cretaceous. Antarctica might have served here as a connection as it was still within close reach of both continents after the Gondwana breakup (McLoughlin 2001 and references therein). During that time, the Antarctic landmass had a warm and humid climate (Dingle and Lavelle 1998). Further evidence of the evolution of Viscaceae in the Upper Cretaceaous comes from the Cretaceous Gondwana origin and subsequent diversification of modern frugivorous birds (the most important seed dispersers of *Viscum*), for which the Antarctic also represents an important factor for the distribution pattern of extant birds (Cracraft 2001). The *Dendrophthora/Phoradendron* clade may have evolved by vicariance as a consequence of a general temperature drop around the Cretaceous/Tertiary boundary. Critical in this respect will be the genus *Arceuthobium* because its species occur in both the Old and New World. However, the selection of *Arceuthobium* species for our study is not adequate to shed any further light onto this issue. We would also like to point out that our Likelihood and Bayesian inference place *Ginalloa* (South-East Asia) as sister to *Dendrophthora*/*Phoradendron* (New World), which differs from earlier, maximum parsimony and ML analysis, where it is sister to *Korthalsella* (South East Asia, Australia, Indian and Pacific Islands).

Puzzling is the reconstructed Late Jurassic/Early Cretaceous origin of many of the included Santalales families, even though some of the fossils used herein were also taken for primary calibrations earlier (e.g., Vidal-Russell and Nickrent 2008). The stem (root) age is in congruence with angiosperm-wide studies of Bell et al. (2005), Naumann et al. 2013, or Magallón et al. (2015), whose representations of all flowering plant lineages came at the expense of a meager sampling of the Santalales diversity. However, our family-based age estimates are partly divergent from earlier reports. For example, with 81–68 mya, the origin of Viscaceae was inferred as much younger by Vidal-Russell and Nickrent (2008), who included only two species of *Viscum* and one of *Arceuthobium*. In contrast, our estimation of the crown group age of Loranthaceae (68.24–42.31 my) are close to or even congruent with earlier studies (e.g., Grímsson et al. 2017; Liu et al. 2018). The focus and, thus, sampling strategies and tree topologies differ largely between earlier works and ours, and so did the preferred molecular dating approaches regarding the employed methods, software programs, models, type and number of genetic markers, fossil information, calibration and constraint strategies. The main difference between ours and comparable existing work is that we refrained from applying any sort of root age constraint or other maximum age constraints. We believe that analysis restrictions based on earlier inferences – no matter how meticulously these were carried out – bear the risk of amplifying error, and that fossil records can not provide evidence of a maximum age. Hence, we present here an alternative age hypothesis for Viscaceae and prefer to avoid judging the correctness of our results versus those of others. To test existing (and newly emerging) hypotheses of divergence times in groups of the sandalwood order, a comprehensive evaluation of an order-wide approach that also assesses the effect of topological uncertainties and methodological robustness is desirable.

Our ancestral area reconstruction does not assign the *Viscum* ancestor unequivocally to one area, but assumes a widespread range comprising Africa, and most probably, Australasia, and Madagascar. Starting from this widely-distributed ancestor, the crown group of *Viscum* seems to have evolved during the early Eocene. The colonization of Australasia and Madagascar by *Viscum*, therefore, must have occurred via long-distance dispersal, assuming the most likely scenario of an African origin of the stem group is inferred correctly. Within clade A, *Viscum* reached continental Asia in the Oligocene from Africa, potentially via India (Fig. 2 and Fig. 3), which is in line with *Viscum* fossils from the Miocene flora of Eastern Georgia (Dzhaparidze 1979). From continental Asia, *Viscum* dispersed to North-Saharan Africa and Europe. The Madagascan clades C and E, together with the Australasian clade D separated in the Eocene from the remaining African clades and the second Asian clade I. Within clade E, the African continent was re-colonized from Madagascar twice, probably through intra-African bird migration. A vicariant event about 30 million years ago separated the two lineages of the Madagascan clade E and the Australasian clade D, which contains two species endemic to Australia. *Viscum* most likely spread to Australasia again between 24 (LN) and 19 (CIR) mya from Africa (clade I) via long distance dispersal. At ca. 25–20 mya (Oligocene), when said split seems to have occurred (Fig. 2), Africa was well separated from Madagascar and Australia, not contiguous with India (and then Australia) since the Cretaceous. Thus, vicariance from direct land mass connections seems unlikely. Stepping stone dispersal via the Kerguelen Plateau, Crozet Plateau, and Ninetyeast ridge represents an alternative explanation, one that has also been proposed for some palms and birds, the latter of which important seed dispersers of *Viscum* (Carpenter et al. 2010; Liu et al. 2018). With some certainty, continental Asia was colonized by *Viscum* at least three times independently from Australasia. In the light of our data, we cannot exclude the possibility of colonizing Australasia outward from Africa via continental Asia. There are well known flyways of migrating birds connecting Africa and continental Asia, while there are no such connections known between Africa and Australia. However, flotsam carrying or infected with fruiting mistletoes, whose berries might get picked up and placed on a new suitable tree, represents another alternative to the lack of flyway routes. Dispersal within the Asian and Australasian regions could be caused by birds migrating along the East-Asian-Australasian flyway, using again islands as stepping stones. Within the African continent, the present-day diversity hotspot of *Viscum*, different clades evolved during the Oligocene with a hotspot in the South-African Cape region (Clades F, G, H). The most recent African clade evolved in the Miocene with a migration towards the Southeast and east of Africa, maybe aided by the establishment of more open woodlands and savannahs during the early Miocene (Jacobs 2004).

### 4.3 Multiple independent reductions of leaves and switches of the reproductive system

The reduction of leaves evolved at least ten times independently in eight of the here defined ten clades (Fig. 2), thus confirming the observations of Danser (1941). At least some species in most geographical regions (Africa, continental Asia, Australasia, and Madagascar) possess scale leaves. Moreover, there is a trend towards this habit prevalent in Africa, and all the species contained in the youngest African clade J possess only vestigial leaves. This drastic reduction in photosynthetic leaf tissue might be a contributor to an advanced parasitic habit for viscaceous mistletoes in general.

Many *Viscum* species are dioecious, reflecting the general trend of a slight overrepresentation of dioecy in parasitic plants compared to nonparasitic angiosperms (Bellot and Renner 2013). Lacking knowledge of the phylogenetic relationships within the genus, the predominance of monoecy within the remaining genera of Viscaceae led Wiens and Barlow (1979) to assume that *Viscum* had evolved from a monoecious ancestor. Our analyses that also considered alternative tree topologies (Fig. 2; Fig. S6 and Table S7) clearly indicate that dioecy is the ancestral reproductive mode in *Viscum*. Despite the higher risk of pollination failure that unisexuality brings, particularly where population densities are small, dioecy guarantees the exchange of genetic material derived from genetically distinct individuals (assuming there is no sex switching), among others by eliminating the opportunity for selfing (as present in bisexuals). Efficient gene flow within and between populations could thus be a beneficial driver in the parasite-host arms race, allowing the parasite to rapidly overcome evolving resistances in host populations or to evolve new mechanisms to exploit novel host systems. However, unisexual plants cannot establish new populations successfully in the absence of the opposite sex. Successful mating in dioecious species inevitably requires two individuals of the opposite sex, plus pollination agents, to overcome the distance between both. The same may apply for shifts in the parasite’s host range. In addition, a seed-shadow handicap (i.e., male plants do not contribute to seed dispersal, through which fewer seeds of dioecious populations may reach new local patches than hermaphrodite ones; Heilbuth et al. 2007) may contribute to the trend towards monoecy, which we find to have evolved at least eight times independently. Therefore, the evolution of this form of hermaphroditism as seen multiple times within *Viscum* also represents an evolutionary advantage for (long-distance) dispersal events – despite the risk of inbreeding-depression. Thus, the ability to self-fertilize apparently may have aided the dispersal and subsequent successful establishment of monoecious *Viscum* species. It is noteworthy to mention that also the dioecious *Viscum album* occurs over great geographic distances. However, the evolutionary success and dispersal range of the European mistletoe may be linked primarily to its broad host range, unparalleled within the genus *Viscum* (Fig. 4; Table S5). The ability to parasitize a great diversity of plants appears to be a rather derived character, which has evolved independently. There is little evidence from our analysis that a direct link exists between the independent reduction of leaves or the shift in the reproductive mode with shifts in host ranges or increasing host specificity in *Viscum* (Figs. 2 and 4). While no causal relationship between reproductive traits and host selection is evident, the reduction of leaves to minute scales, which *de facto* means the loss of photosynthetic surface, may indicate an increasing parasitic specialization and dependence upon host-derived photosynthates. The extreme reduction of surface area for photosynthesis, which in *Viscum* can be reduced to photosynthetic stems only, and thus autotrophic carbon supply, in consequence to this near-loss of leaves requires that the uptake of organic carbon is secured, perhaps through a more efficient connection to its host’s vascular tissue. Leafy mistletoes such as *V. album* obtain a large though host-specific fraction of carbon from their host although they establish only xylem connections (Richter and Popp 1992).

*Viscum*, as well as other obligate parasites, depend on the availability of a compatible host plant, which adds an additional isolating barrier during the speciation process that is absent from nonparasites. Beyond that, geographic barriers are important, too, not only for the mostly bird-driven distribution of the parasites but also for the host. The sedentary nature of plants, thus, may contribute a plausible explanation why many parasitic plants tend to have broader host ranges compared to most animal parasites, whereas the specialization on only a few closely related host species (as in *V. minimum*) is observed rather rarely. Thus, the interaction of factors like the degree of host specificity, the abundance of and distances between different host individuals, plus the abundance of dispersing birds and the distance these dispersers cover may also have played a critical role during the evolution of the genus *Viscum* and mistletoes in general.

To summarize, we have shown here that morphology, geographic range, and the similarity in host preferences are inconclusive indicators of phylogenetic relatedness in *Viscum* mistletoes. Despite the probable beneficial switch of mating systems within this genus aiding their establishment in new geographic areas, the biotic-abiotic factors and their interactions driving the speciation and diversification within *Viscum* remain to be clarified, thus leaving us with the pressing question whether mistletoes might have diversified in the shadow of birds.

## 5. Acknowledgements

The authors thank the Herbarium of the Natural History Museum Stuttgart (STU), Australian National Herbarium (CANB), The Botanical Gardens of Bonn, Herbarium of the University of Bonn (BONN), The National Herbarium of the Netherlands (WAG), The Herbarium of the Missouri Botanical Garden (MO), Herbarium of the Real Jardín Botánico Madrid (MA) for providing specimens and their permissions for destructive sampling. We appreciate the efforts of three anonymous Reviewers and the editorial team and thank them for their critical and insightful comments on an earlier version of this manuscript. This research was supported by a grant from the GIF, the German-Israeli Foundation for Scientific Research and Development (G-2415–413.13 to S.W.), in addition to intramural funds from the Universities of Muenster (S.W., K.F.M.) and Bonn (D.Q.). S.W. is a fellow of the *Emmy Noether*-program of the German Science Foundation (DFG, WI4507/3–1).

## 6. Authors’ contributions

K.M. and S.W. designed this study, generated and analyzed the data, and wrote the manuscript. D.L.N. and K.F.M. contributed to the study design, to performing this research, and to critically revising the manuscript. M.K. and D.Q. contributed to the phylogenetic analyses or data generation. All authors have read and approved the final manuscript.

## 10. Supporting Information

Figure S1 Topological constraints inferred with the four-marker data set 1 used as topological constraints for ingroup inferences^1^

Figure S2 Phylogenetic trees obtained from BI and ML analysis of the 110-taxa data set 2 used as input data for ancestral state estimations^1^

Figure S3 Phylogenetic tree and summary of topological conflicts within the genus *Viscum* and outgroups^1^

Figure S4 Cladograms with support values from alignment optimizations of dataset 2^1^

Figure S5 Phylograms obtained from alignment optimizations of data set 2^1^

Figure S6 Node delimitation for PhyloBayes and BayesTraits results, as detailed in Table S7 Table S1 Voucher information and NCBI Genbank accession numbers of all taxa used for the reconstruction of a backbone guide tree with data set 1

Table S2 Voucher information and NCBI Genbank accession numbers for all taxa used for the reconstruction of the final 110-taxa phylogeny with data set 2

Table S3 Geographic ranges for ancestral area estimation^2^

Table S4 Morphological features for ancestral state estimation^2^

Table S5 Host information for *Viscum* and outgroup taxa^2^

Table S6 Results of the *LaGrange* analyses

Table S7 Results of the *PhyloBayes* and *BayesTraits* analyses

## Footnotes

1 Phylogenetic trees also available in *newick* format from *Mendeley Data*.

2 Trait data also available in excel table format and as easy-to-parse plain text files (tab-delimited with comment lines) from *Mendeley Data*.

